# Mechanistic target of rapamycin/blood-testes barrier mechanism mediates acceleration of sperm epigenetic aging by environmental factors

**DOI:** 10.1101/2025.02.25.640214

**Authors:** Olatunbosun Arowolo, Oladele A Oluwayiose, Jiahui Zhu, Oleg Sergeyev, Emily Houle, J Richard Pilsner, Alexander Suvorov

## Abstract

Our previous research suggested that mechanistic target of rapamycin (mTOR)/blood-testis barrier (BTB) mechanism is involved in the regulation of the rates of epigenetic aging of sperm, where increased activity of mTORC1 opens BTB and accelerates epigenetic aging and increased activity of mTORC2 produces opposite results – increases BTB integrity and rejuvenates sperm epigenome. In the present study, we use our newly developed epigenetic clock model to investigate whether the mTOR/BTB mechanism is involved in the epigenetic reprogramming of sperm in mice exposed to heat stress (HS) and cadmium (Cd). Our findings show that both mTOR-dependent BTB disruption caused by HS and mTOR-independent BTB disruption due to Cd exposure accelerate sperm epigenetic aging, resulting in similar changes to sperm DNA methylation patterns. These results suggest that the mTOR/BTB mechanism is a novel molecular pathway through which environmental stressors influence sperm epigenetic aging, and this pathway may be relevant to a broad range of factors, including environmental, lifestyle, dietary, and health influences.

## Introduction

Research on sperm epigenetics over the last two decades demonstrated that DNA methylation patterns in spermatozoa may be affected by a broad range of environmental and lifestyle factors, health conditions, and aging. For example, it was shown that diet [1, 2], physical exercise [3, 4], alcohol consumption [5], psychological stress [6, 7], aging [8, 9], smoking [10, 11], and exposures to environmental toxicants [12–16] all have effects on DNA methylation profiles of sperm.

It is long known that after fertilization, paternal DNA methylation undergoes reprogramming to erase parental epigenetic marks and establish totipotency in the developing embryo. This understanding was modified in recent years, as according to emerging research, many DNA methylation marks may survive reprograming in early embryogenesis and affect embryo development [17–23]. The role of sperm epigenetics in the transfer of inheritable information to the next generation was suggested by many lines of population research. For example, epidemiological studies demonstrated that advanced paternal age increases risks of poor pregnancy outcomes [24, 25], and adverse health of the offspring [26–28]. Offspring of advanced age fathers have higher odds for the development of early cancer [29]; higher incidence of schizophrenia [30] and autism [27, 28, 31, 32] among other conditions. In laboratory rodents advanced paternal age resulted in shorter lifespan [19], altered vocal communication [33], and reduced exploratory behavior [18] in offspring. Other paternal factors also exerted intergenerational effects in laboratory animals. For example, paternal psychological stress affected risk-taking behavior [17], and many paternal factors, including psychological stress [17], prediabetes [34], and in utero undernourishment [35] resulted in metabolic dysfunction in mouse offspring. Taken together this evidence suggests that the sperm methylome is a significant channel for the transfer of inheritable information to the next generation [36].

A recent comprehensive review of age-dependent changes in mammalian sperm demonstrated that aging is the most powerful known factor associated with sperm epigenomics patterns [37]. Aging produces profound changes in sperm epigenomes of laboratory models [38–40] and humans [8, 9] and affects all known epigenetic mechanisms, including DNA methylation [40–42], small non-coding RNA [43–46] and histone modification [47]. For example, we identified 5,319 age-dependent differentially methylated regions (DMRs) [42] and 1,384 differentially expressed small non-coding RNA [43] between 65 and 120-day-old rats. Additionally, published data including ours [19, 42, 48] show that genes associated with age-dependent DMRs or targeted by age-dependent sncRNAs are enriched with major developmental pathways, including nervous system-related signaling, Wnt, Hippo, mTOR, and Igf1 [19, 33, 42, 44–46, 48]. Likewise, research in two independent cohorts demonstrated profound associations between age and sperm DNA methylation [49, 50].

Changes in sperm DNA methylation induced by age or stressors often follow common patterns, which cannot be explained by random causes [37, 51]. Specifically, many studies exploring effects of various factors on sperm DNA methylation, including age, diet, chemical exposure, heat stress, and psychological stress, report that differentially methylated regions (DMRs) are associated with genes enriched for similar biological categories, such as metabolism, embryonic development, neurodevelopment, and behavior [19, 20, 22, 34, 35, 40, 42, 50, 52–59]. The profound effect of aging on sperm DNA methylation, taken together with similarity of effects of different factors on sperm epigenome, suggests that environmental and other factors may exert their effects on sperm methylome via acceleration or deceleration of the natural age-dependent changes in the sperm methylome. Indeed, urinary phthalate metabolites in humans and exposure to brominated flame retardant BDE-47 in rats accelerated the rates of aging of sperm methylome [43, 60, 61].

Our recent study shed light on molecular mechanisms involved in the regulation of aging of sperm methylome [40]. Using transgenic mice, we demonstrated that epigenetic aging of sperm can be accelerated or decelerated by experimental shifts of the balance of the two complexes of the mechanistic target of rapamycin (mTOR) – serin-threonine protein kinase, a molecular hub for many intercellular pathways expressed at highest levels in testes in comparison to other tissues in humans [62] and mice [63]. In our experiments, Sertoli-specific suppression of mTOR complex 1 (mTORC1) resulted in acceleration of epigenetic aging of sperm, while suppression of mTOR complex 2 (mTORC2) rejuvenated sperm methylome. Interestingly, the same mechanism regulates permeability of the blood-testes barrier (BTB) – the shift of the balance of the two complexes in Sertoli cells in favor of mTORC1 opens the BTB, while the shift in favor of mTORC2 promotes the BTB integrity. The BTB is the tightest blood tissue barrier [64, 65] that regulates the biochemical environment in the lumen of the seminiferous epithelium where germ cells undergo epigenetic reprograming [66–68]. Animal studies show that age-dependent deterioration of the BTB integrity is the cause of fertility declines with age [69–71]. These observations suggest that mTOR effects on epigenetic aging of sperm may be mediated by changes in the BTB integrity.

Thus, in this study, we hypothesize that environmental factors affecting mTOR and/or BTB affect the rates of epigenetic aging of sperm. To estimate aging acceleration or deceleration, we developed a mouse sperm epigenetic clock model. The first epigenetic clocks were developed in 2013 as statistical models linking the levels of methylation of selected DNA regions with chronological age in humans [72, 73]. Many modifications of epigenetic clocks were developed since then using different cell and tissue types, model organisms and using different training approaches (e.g. health status instead of chronological age or combination of both) [74–76]. Application of these models demonstrated that epigenetic age acceleration is associated with early menopause [77], cancer, cardiovascular and all-cause mortality [78], allergy and asthma in children [79], smoking [80], exposure to air pollution [81–83], childhood exposure to violence [84], and others. It was shown, however, that epigenetic clock models trained on somatic tissues do not have predictive relevance to sperm [85]. Several versions of sperm epigenetic clocks were built for humans [86–88], however, a sperm epigenic clock has not been developed for mice to our knowledge. Thus, in the current study, we fill this methodological gap and develop murine sperm clock model to estimate epigenetic age shift by environmental factors.

In this study, we use heat stress (HS) as a model of potentially mTOR-dependent BTB disruption by environmental factors. HS has been shown to activate mTORC1 signaling in skeletal muscles in humans [89] and rats [90] and in turkey muscle stem cells [91, 92]. HS also induced the disruption and leakage of the BTB [93, 94] and reduced expression of structural BTB molecules (connexin-43, ZO-1, vimentin, claudin-1, claudin-5) in Sertoli cells [95]. Similarly, expression of tight junction components (occludin, claudin-3, and ZO-1), was reduced and localization of occludin and ZO-1 was lost from the BTB after a mild scrotal heat exposure in mice [96] resulting in BTB degradation which required 10 days to recover [96]. Expression of occludin and its localization to BTB was also decreased in boars given local scrotal exposure to 42°C for 1h [97].

Analysis of heat stress effects on sperm epigenetics gained importance in recent years due to the climate change, which according to the WHO, is the single biggest health threat facing humanity [98]. One way climate change impacts health is via increasing frequency of heat waves worldwide [99]. For example, marine heatwaves are twice as likely as they were in 1980 [99]. It is predicted that an event that would occur every ten years, would occur every other year if global warming reaches 2°C [100]. According to the US EPA, heat is the leading weather-related killer in the US [101]. Although the testis is recognized as the tissue most susceptible to HS [94] consequences of regular exposure to HS on sperm epigenome remain obscure. A heatwave is defined as five or more consecutive days of prolonged heat with a daily maximum temperature higher than the average maximum temperature by 5°C or more [102]. Given that many humans have an opportunity to avoid prolonged exposures to HS during heat waves using air-conditioned environment and other means, we developed an environmentally relevant short-term acute intermittent whole-body HS exposure protocol to simulate human exposures. In this protocol mice are subjected to alternating periods of HS exposure for 20 min per day, for 5 consecutive days with 2 weeks recovery periods.

To model environmental disruption of the BTB via mTOR independent mechanism, we used cadmium (Cd) exposure. Cd is a well-recognized BTB disruptor [103–105] which affects BTB in an mTOR-independent manner [106, 107]. Mechanistically, Cd disrupts the BTB by up-regulating transforming growth factor β3 (TGF-β3), which then activates p38 MAPK signaling, and downregulates FAK protein expression, which regulates tight junction proteins such as ZO-1 and occludin [106–110]. Cd is also known to be a potent modifier of sperm DNA methylation in mice [111]. Although Cd toxicity is not a focus of the current application per se, our experiments may contribute to better understanding of an important public health issue as millions of people worldwide are chronically exposed to this metal [112].

Thus, in the current study, we tested a hypothesis that potentially mTOR-dependent (HS) and mTOR-independent (Cd) disruption of the BTB by environmental factors accelerate epigenetic aging of sperm in mice. We demonstrate epigenetic age acceleration by 31.7 and 42.3 days by the two factors respectively. This study suggests that mTOR/BTB mechanism is a common mechanism converting paternal exposures to acceleration of their sperm epigenetic aging. This study also provides murine sperm-specific epigenetic clock model, which can be used in various future applications.

## Materials and Method

### Animals and Treatment

C57BL6 male mice of different age were purchased form the Jackson Laboratories and used for sperm epigenetic clock model development or for exposure experiments. All animals were allowed to acclimate for one week and were single-housed in a temperature (23 ± 2°C), and humidity (40 ± 10 %) controlled environment, with a 12-h light/dark cycle, and food and water available ad libitum. For sperm epigenetic clock, 42 animals were euthanized at 14 different ages as follows (postnatal day – number of mice): 69 - 4, 80 - 4, 107 - 4, 118 - 3, 165 - 2, 186 - 1, 190 - 4, 225 - 3, 252 - 1, 265 - 4, 276 - 4, 300 - 1, 330 - 4, and 334 - 4. For exposure experiments, twenty 100-days old mice were assigned to one of the following groups (n = 5 per group): control group, 31.5°C and 34.5°C HS groups (31.5°C and 34.5°C groups respectively), and cadmium group (Cd).

Cd group was exposed daily to 1 ul/g body weight of water solution of CdCl_2_ (Cat. # 202908, analytical grade, Millipore Sigma) to achieve a dose of 2 mg/kg of body weight per day. A similar exposure dose was used in a previous study demonstrating significant changes in sperm DNA methylation [111]. Exposure to Cd solution was conducted by feeding from a tip of pipettor, a method validated in our previous research [43, 113–115]. Control and HS groups were exposed same way to 1 ul/g body weight of drinking water. All exposures were done for 78 days to ensure coverage of two spermatogenesis cycles.

HS mice (31.5°C and 34.5°C groups) were exposed to elevated temperatures using infrared ceramic heat emitters [116] for five days, for a duration of 20 min per day, to mimic human intermittent exposure to ambient and conditioned environment during heat waves. Five-day periods of heat stress alternated with two-week recovery periods for the duration of the experiment. Thus, over 78-day protocol, every animal was subjected to four complete treatment periods (5-day HS + 14 days recovery) followed by 2 days of HS. Previous research demonstrated that mice can tolerate continuous exposure to 33–35 °C for several days although it alters animal physiology [116], while exposures to 40-50°C for 0.5-2 hours results in the death of a large number of laboratory animals [117–119]. The HS exposure device consisted of a plastic cage with two ceramic hit emitters (BOEESPAT Ceramic Heat Emitter 100W) installed above the cage and controlled by a digital temperature controller (XIEHUZA Backlit Digital Temperature Controller) with its sensor installed at the bottom of the cage. Five additional thermometers were installed in the center and each corner of the cage to control for even distribution of heat in the cage. Infrared heat lamps are safe as emitters do not produce any type of light to which murine retinal cells are sensitive [117].

Health parameters of all animals were monitored daily for the duration of experiment, and all animals were euthanized by cervical dislocation on 78^th^ day of exposure. At euthanasia, epididymal sperm and testes were collected from every animal. Testes samples were snap frozen in liquid nitrogen and stored at -80°C. All procedures followed the guidelines of the National Institutes of Health Guide for the Care and Use of Laboratory Animals and the approval for this study was received from the Institutional Animal Care and Use Committee at University of Massachusetts, Amherst.

### Sperm collection and DNA extraction

Caudal epididymal sperm was collected using procedure described in our previous research [14]. In short, epididymides were incised, cut three times, and incubated at 37°C for 30 minutes in 1 ml of sperm wash buffer (Cat. # ART1006, Origio, Denmark). After incubation the epididymides were removed, and sperm samples were used for DNA extraction following the rapid method developed in J.R. Pilsner’s laboratory [120]. In short, sperm samples were subjected to a one-step density gradient centrifugation over 40% Pureception buffer (CooperSurgical Cat # ART-2100) at 600 g for 30 min in order to remove possible somatic contamination. Sperm cells were homogenized with 0.2 mm steel beads at room temperature in a mixture containing guanidine thiocyanate lysis buffer and 50 mM tris(2-carboxyethyl) phosphine (TCEP; Pierce, Rockford, IL, USA). The lysates were column purified using QiaAMP DNA mini-Kit (Qiagen, Cat # 56304) and DNA quality of the eluate was determined using Nanodrop 2000 Spectrophotometer (Thermo Scientific, Somerset, NJ). The DNA samples were stored at -80°C.

### Infinium array analysis and results processing

Sperm DNA methylation analysis was conducted for all mice in control, 34.5°C, and Cd groups in the genomic facility of Wayne State University using Infinium mouse Methylation Beadchip (Illumina, San Diego, CA, USA) array with a genome-wide coverage of 285K methylation sites. In the bead chip array, each CpG methylation level was estimated based on the intensity of unmethylated and methylated probes. A total of 288,658 probes were identified and were preprocessed using the sesame function which included background and dye-bias corrections as well as the removal of SNPs and repeated probes in R. The minfi package was used to remove probes that are below background fluorescence levels, adjust the difference in Type I and II probes, and correct for technical variations in the background signals [121]. The ComBat function was used to correct batch effects [122]. This resulted in a total of 254,410 CpGs for downstream analyses. The results were reported in beta values, which is a measure of methylation levels from 0 to 1. The beta values are converted to M-values by log2 transformation, as this has been reported to be a more valid approach for differential methylation analysis and better conforms to the linear model homoscedasticity [123]. DMR were identified using the A-clustering algorithm [124].

### Epigenetic clock model

A CpG-based sperm epigenetic clock was developed using methylation data on 254,410 CpGs. Methylation data were fed directly into the elastic net algorithm, which selected 64 CpGs for the final clock model. These 64 CpGs are from a total of 42 samples. The 42 samples used for the clock were then split into two subsets (20/80%) for cross-validation (n=9 for 20%, and n=33 for 80%). The elastic net is a penalized regression model that combines predictions from two algorithms, LASSO, and ridge [125]. The final clock was used to predict the age of the sperm of the control and the treated groups.

### Enrichment analysis

To analyze biological categories enriched by genes associated with the 64 CpGs used for building the clock, we first identified coordinates of these CpGs using the Illumina manifest. Annotation to reference genome mm10 was done using default settings of the Annotatr package in R. The nearest genes were identified with transcription start site (TSS) 1500 using GRCh37 assembly data from ENSEMBL via the annotatePeakInBatch function from the ChIPpeakAnno package. To identify changes in DNA methylation induced by Cd exposure and HS exposure DMRs were identified using A-clustering algorithm [124] in comparison with controls. Same approach was used to identify biological categories enriched with regions undergoing methylation change with aging. For this later analysis we compared DNA methylation in sperm of young mice (combined group of four 80-days old and four 107-days old animals) and “old” mice (combined group of four 330-days old and four 334-days old animals). The lists of genes associated with DMRs for aging and environmental factors and CpGs for epigenetic clock were submitted to Metascape to identify enriched biological categories [126]. Additionally, we conducted enrichment analysis for 500 randomly selected CpG probes to test if enrichment analysis is biased towards ontology terms overrepresented in gene annotations. Finally, we used Gene Set Enrichment Analysis (GSEA) [127] to visualize changes in methylation induced by aging and environmental factors.

### mTOR activity in testes

To assess changes in mTOR pathway, we measured the phosphorylation of well-characterized mTORC1 and mTORC2 targets, phospho-p70S6 kinase (Thr389) and phospho-Akt (ser473), respectively, in homogenates of whole testes using ELISA (ab176651 and ab253299 respectively, Abcam) in accordance with protocols recommended by the manufacturer. Concentration of phosphorylated proteins was expressed as relative quantification with average value for control group set to one.

### Statistical analysis

Shapiro-Wilk normality test was first done to assess if our data have normal distributions. Differences in mTOR complexes activity between control and treatment groups were than identified using T-test; and differences in epigenetic age shift between control and treatment groups were identified using Welch’s T-test, as variances for the latter were not equal between groups. We considered p-value ≤ 0.05 as statistically significant difference. The statistical approaches for bioinformatic analyses are described in corresponding sections above.

## Results

Body weights measured the last day of treatment were as follows (mean ± SE): 29.5 ± 0.9, 31.1 ± 0.2, 28.7 ± 1.5, and 27.8 ± 1.1 for control, 31.5°C, 34.5°C, and cadmium group respectively. Body weights were not significantly different in all treated groups as compared with controls. Exposures to environmental stressors reduced testis weights. Testis weight was 81.2±3.5 (mean±SE) in control animals while in treated mice it was 78.5±2.3 in 31.5°C group (*p*=0.3), 70.4±4.1 in 34.5°C group (*p*= 0.04), and 61.8±2.0 in Cd group (*p* = 0.0005) (Fig. 1A). No changes in outwardly behavior and appearance were observed in treated animals as compared with controls except increased motor activity in both HS groups.

**Figure 1.**
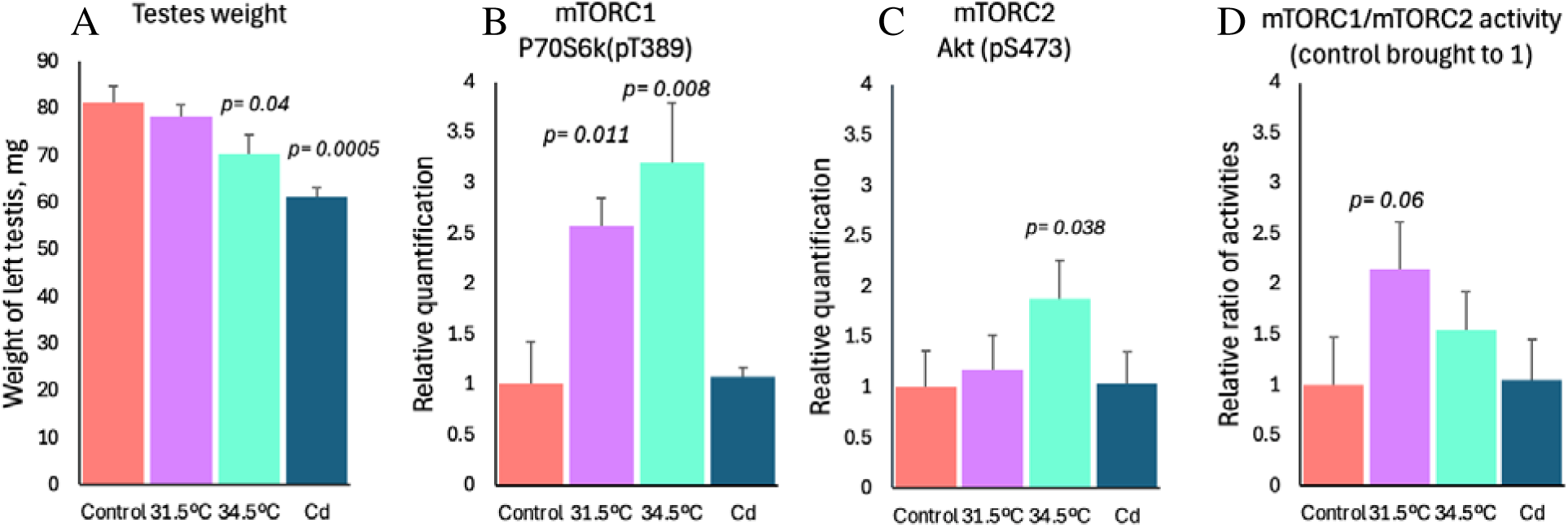
Heat stress and Cd induced reduction of testes weight (A) and changes in activity of mTOR complexes in testes homogenates: (B) activity of mTORC1, (C) activity of mTORC2, and (D) ratio of mTORC1/mTORC2 activities. mTOR complexes activity was measured by ELISA for phospho-proteins p70S6K and Akt – readouts of mTORC1 and mTORC2 activities respectively. P-values are for t-test comparison with controls.

### Heat stress but not cadmium exposure induced mTOR activity in testes

We analyzed activity of mTOR complexes in testes homogenates by measuring relative concentrations of phospho-proteins p70S6K and Akt – readouts of mTORC1 and mTORC2 activities respectively (Fig. 1B-C). Our result showed that HS, but not Cd, affects mTOR activities. HS at both temperatures of 31.5°C and 34.5°C significantly induced mTORC1 activity around 2.5 and 3-fold, respectively. Changes of mTORC2 was non-significant at 31.5°C but reached a significant 1.9-fold activation at 34.5°C. In addition, our results also showed that HS, but not Cd, affects the balance of mTOR complexes, such that both HS at 31.5°C and 34.5°C resulted in the shift of the balance of mTOR complexes in favor of mTORC1 around 2- and 1.5-fold, respectively (Fig. 1D). This shift was non-significant at 34.5°C and semi-significant (p = 0.06) at 31.5°C.

### DNA methylation changes induced by aging, HS, and Cd affect similar biological categories

To determine the biological significance of sperm DNA methylation changes due to 34.5°C HS and Cd exposure, we identified genes associated with differential methylation (DMRs) between each exposure, and control (Supplemental Table 1). We also identified genes associated with DMRs between young and old mice. The lists of genes associated with significant DMRs induced by aging, HS and Cd exposures are enriched with similar age-dependent developmental categories (Fig. 2). Common enriched categories include cellular development (cell-cell adhesion, cell junction organization and morphogenesis), neuronal development, and axon guidance. To test if similarity between enriched categories is due to a bias towards ontology terms overrepresented in gene annotations, we also conducted enrichment analysis for 500 randomly selected CpG probes. This analysis shows that similarities of effects of aging, Cd exposure, and HS on categories enriched with differentially methylated CpGs cannot be explained by annotations bias.

**Figure 2.**
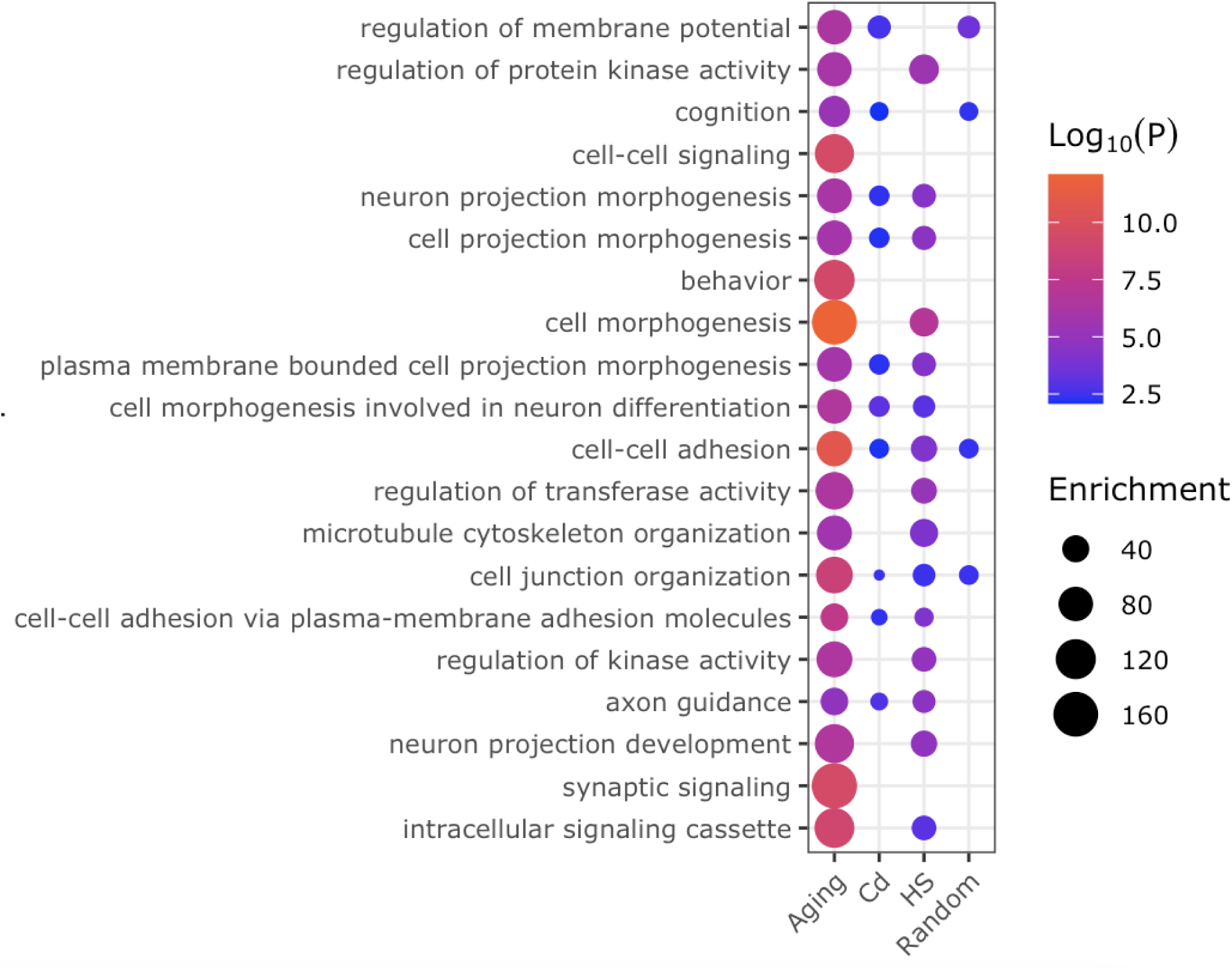
Enriched biological categories due to aging, cadmium exposure, heat stress, and those associated with randomly selected genes.

### Aging and stressors produce similar changes in sperm DNA methylation

According to our hypothesis mTOR-dependent and mTOR-independent disruption of the BTB will change sperm DNA methylation via acceleration of epigenetic aging of sperm. If this hypothesis is true, aging and both stressors (HS and Cd) would produce similar changes in sperm DNA methylation. To test this experimentally we used GSEA to compare age dependent changes in DNA methylation with DNA methylation in significant DMRs induced by 34.5°C HS and Cd exposure. The results of this analyses demonstrate that DNA regions undergoing hypomethylation with age are also hypomethylated in response to Cd and HS (Fig. 3A, B), while DNA regions undergoing hypermethylation with age also gain methylation in response to Cd and HS exposure (Fig. 3C, D). These results indicate that environmental disruptors of the BTB affect sperm DNA methylation in a way that resembles acceleration of epigenetic aging.

**Figure 3.**
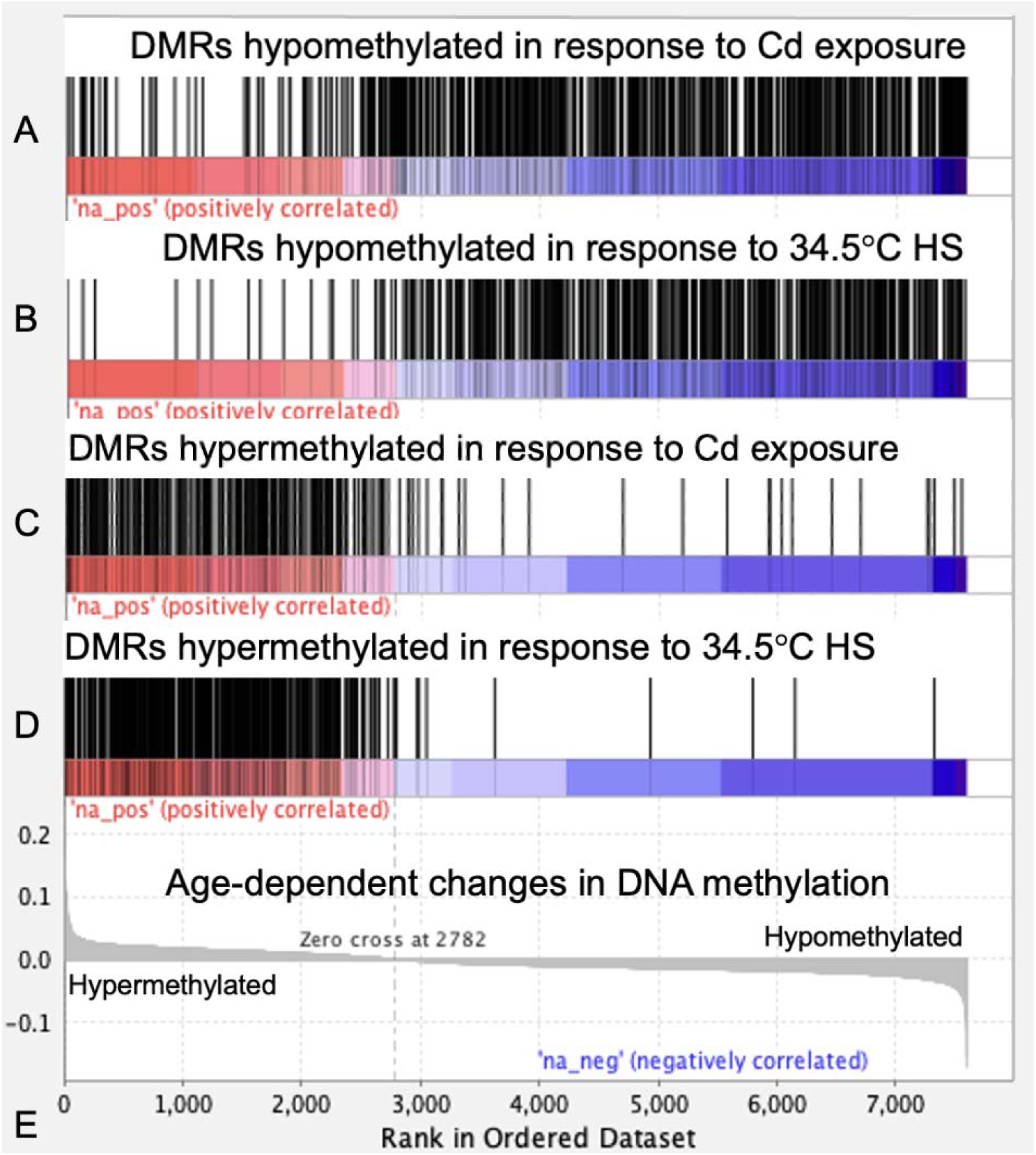
Comparison of hypo- (A, B) and hypermethylated (C, D) DMRs in response to Cd (A, C) and 34.5°C HS (B, D) exposures with methylation changes induced by aging (E) in mouse sperm.

### Mouse sperm epigenetic clock

To analyze shifts in the sperm epigenetic age induced by environmental stressors, we developed a mouse sperm epigenetic clock model (Fig. 4, Supplemental Table 2). Our clock has a predictive capacity with a mean absolute error (MAE) value of 2.3 days for in-sample datasets (Fig. 4A), and 14.3 days for out-of-sample datasets (cross-validation datasets) (Fig. 4B). Genes associated with the 64 CpGs used for sperm clock were enriched for adipocytokine signaling pathway, mRNA metabolic process, learning, and cell morphogenesis (Fig. 4C).

**Figure 4.**
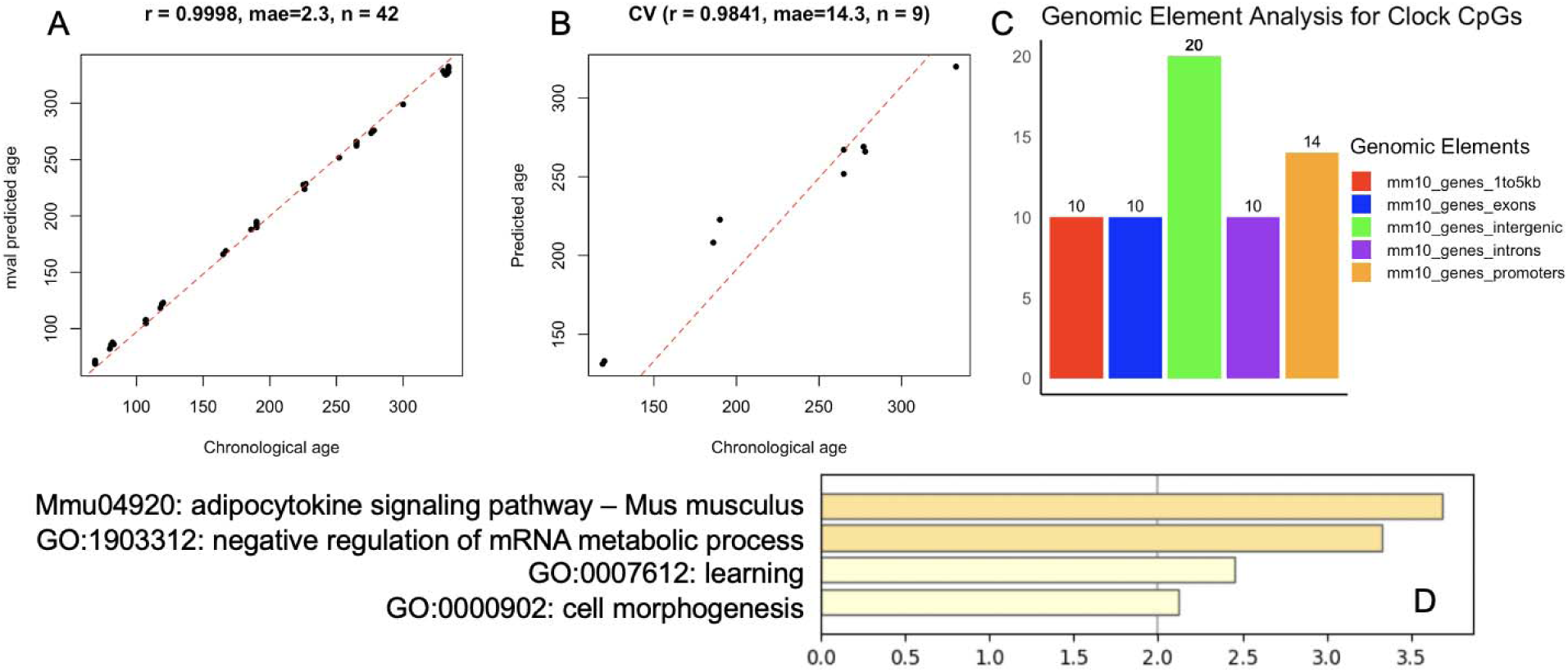
Elastic net selected age-associated sperm methylation for epigenetic clock predicted from linearly correlated CpGs **(A)** Scatterplot of correlation between chronological age and predicted age. (**B)** Scatterplot of correlation between chronological age and out-of-sample, cross-validated (CV) predicted age. **(C)** Overlap of clock CpGs with genomic elements. **(D)** Biological categories enriched with genes associated with 64 clocks CpGs.

### Environmental stressors induce age dependent changes in sperm

In our previous study, we demonstrated that a shift in the balance of mTOR complexes activities in favor of mTORC1 produces changes in sperm DNA methylation matching accelerated aging [40]. Here, we tested if similar shift in mTOR complexes balance in favor of mTORC1 activity by HS will accelerate sperm epigenetic age. Indeed, 34.5°C HS resulted in 31.7 ± 4.5 (mean±SE, *p*=0.0031) days older sperm than same age controls as assessed using our sperm epigenetic clock (Fig. 5).

**Figure 5.**
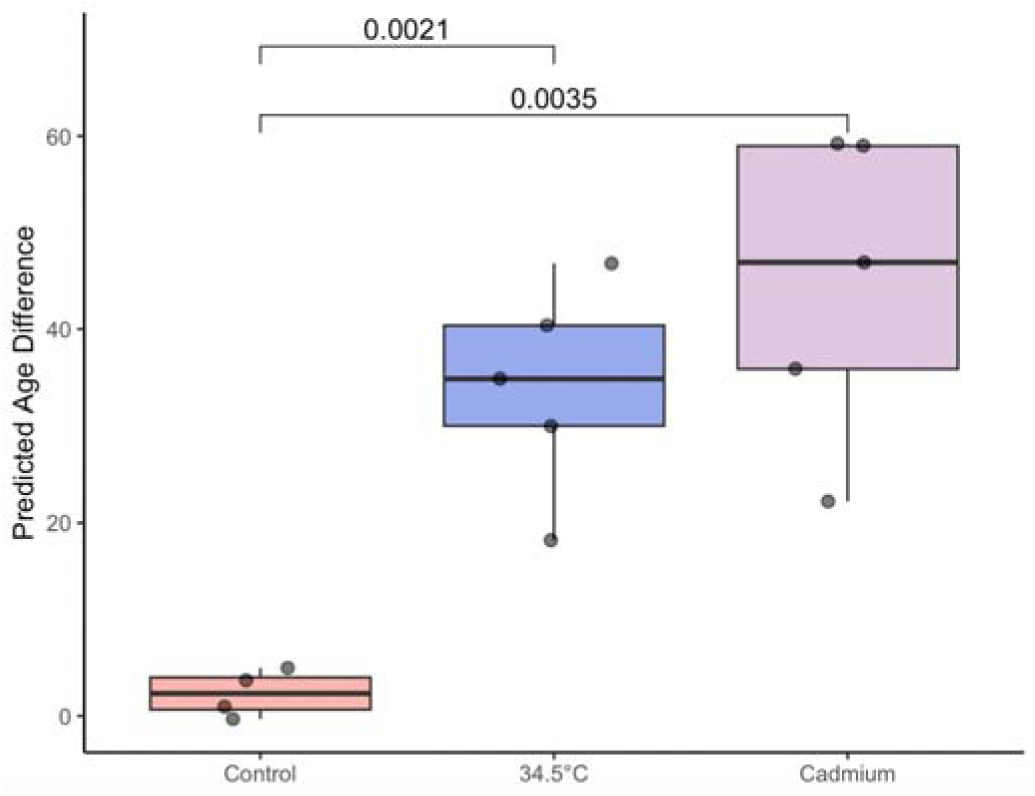
The effects of environmental exposures, Cd and HS, on the sperm epigenetic age (n=4-5/group, p-value are for Welch’s t-test comparison with control group).

Since the balance between the two mTOR complexes in the Sertoli cell regulates the integrity of the BTB [128], it is plausible that the effects of mTOR pathway on sperm epigenome are BTB dependent. To test this hypothesis experimentally we analyzed changes in sperm epigenetic age of mice exposed to Cd, a well characterized mTOR-independent BTB disruptor [129]. Our results support the hypothesis, as epigenetic age of sperm of Cd exposed mice was 42.3 ± 7.2 (mean±SE, *p*=0.0035) days older than epigenetic age of sperm in control mice. (Fig. 5).

## Discussions

Our previous research suggested that mTOR/BTB mechanism is involved in the regulation of the rates of epigenetic aging of sperm, where increased activity of mTORC1 opens BTB and accelerates epigenetic aging and increased activity of mTORC2 produces opposite results – increased integrity of the BTB and sperm epigenome rejuvenation [40]. In the current study, we tested if this mechanism is involved in epigenetic reprogramming of sperm by environmental factors. Specifically, we tested using newly developed epigenetic clock model if putative mTOR-dependent BTB disruption by HS and mTOR-independent BTB disruption by Cd will accelerate epigenetic aging of sperm and will produce similar effects on sperm DNA methylation.

First, we obtained experimental evidence that HS activates mTOR complexes in testes and shifts the balance of their activity in favor of mTORC1, while Cd exposure does not affect mTOR activity. These findings support our hypothesis that HS and Cd are mTOR-dependent and mTOR-independent BTB disruptors respectively. We did not measure the BTB disruption directly as it was well characterized in extensive previous research of HS [93–97, 130–134] and Cd [103–105, 108–110, 129, 135–142] effects on BTB integrity and permeability.

We report that exposures to the reproductive toxin, Cd, and HS, as well as aging produce changes in sperm DNA methylation associated with very similar set of developmental categories. Enrichment of similar developmental categories was also reported with sperm methylation changes in response to a variety of factors including diet, chemical exposure, heat stress, psychological stress, and aging [19, 20, 22, 34, 35, 40, 42, 50, 52–59]. Most intriguing, both environmental factors tested in the current study displayed acceleration of epigenetic aging of sperm. Taken together, these findings suggest that BTB disruptors Cd and HS, along with other previously tested factors, affect sperm DNA methylation by accelerating the natural process of epigenetic aging of sperm.

Our current and recent research provides the following mechanistic model for age-dependent changes in the sperm methylome (Fig. 6). During the reproductive life, the integrity of the BTB is decreasing with age [69–71], resulting in increasingly noisy environment in the apical compartment of the seminiferous epithelium. Events that occur within this compartment include the global methylation drop of around 12-13% in preleptotene spermatocytes and methylation reestablishing during leptotene-pachytene stages [143–145], and changes in methylation of thousands DNA regions during transition from elongating to late spermatids [146]. We suggest that age-dependent increase in the BTB leakage may result in increasingly non-optimal biochemical environment in the seminiferous epithelium and promote stochastic errors in methylation, where regions that are normally hypermethylated in young, untreated animals gain methylation and regions that are normally hypomethylated in young, untreated animals loose methylation [51]. These stochastic events averaged for many spermatozoa and/or organisms form age-dependent semi-linear trends for methylation increase or decrease respectively [40], a phenomenon captured by epigenetic clocks [51, 147]. Importantly, in sperm, some DNA regions are prone to higher stochastic variability in DNA methylation. These regions termed variable methylation regions (VMRs) were identified and characterized in our recent study, which demonstrated that the majority of age-dependent DMRs are also VMRs [51]. In other words, epigenetic aging of sperm is a result of increasing stochasticity of methylation in VMRs.

**Figure 6.**
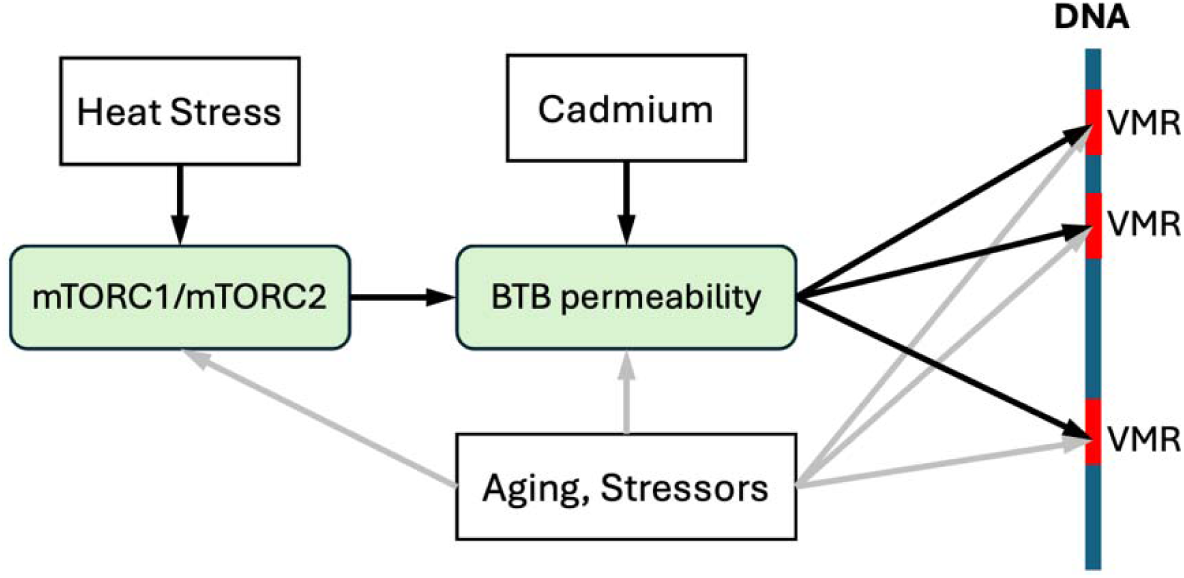
Mechanistic model explaining age/stress dependent changes in sperm DNA methylation. Left to right: heat stress shifts the balance of mTOR complexes activity in Sertoli cells in the favor of mTORC1. This shift in mTOR complexes balance results in increased BTB permeability. BTB permeability may be also affected via mTOR-independent mechanisms, such as Cd exposure. Leaky BTB generates suboptimal biochemical environment in the apical compartment of seminiferous epithelium and increases the rate of stochastic changes in DNA methylation in naturally variable methylation regions (VMRs). Increased stochastic events in VMRs shifts average methylation values in these regions and results in differential methylation. Different nodes of this cascade may be affected by a broad range of stressors and aging.

Findings from this study suggests that HS and Cd increase the rates of stochastic errors in sperm DNA, resulting in accelerated rates of epigenetic aging of sperm. Both stressors are well characterized disruptors of the BTB, and thus, their effects mimic effect of aging on the BTB integrity [69–71]. Given that mTOR pathway contributes to the maintenance of the BTB integrity, a broad range of inputs that merge on mTOR may affect the rates of sperm methylome aging. Similarity of DNA methylation changes induced by various stressors in many published studies [19, 20, 22, 34, 35, 40, 42, 50, 52–59], suggests that they also act via increasing rates of stochastic epigenetic events, although it is unclear if their effects are also mediated via mTOR and/or BTB or other pathways.

This study does not provide a clear understanding of precise molecular mechanisms that translate BTB integrity into sperm DNA methylation changes. As discussed above, age-dependent changes in DNA methylation result from stochastic errors accumulation within sensitive methylation regions [51]. As such, we speculate that increased BTB leakage may generate suboptimal biochemical environment in the apical compartment of seminiferous tubules, which in turn, increases the rate of stochastic errors. The precise sequence of molecular event involved in this causative chain remains unclear and requires additional research.

## Conclusion

We identified the novel molecular mechanism that mediates effects of environmental stressors on sperm epigenetic aging. This mechanism includes mTOR-dependent or mTOR-independent disruption of the BTB integrity. We hypothesize that leaky BTB accelerates sperm epigenetic aging by creating suboptimal biochemical environment in the apical compartment of seminiferous tubules which promotes increased rates of stochastic errors in DNA methylation. Additionally, this research also provides novel mouse sperm epigenetic clock which may be used in a broad range of future studies of sperm DNA methylation utilizing murine model.

## Supplemental Materials

Supplemental Table 1. Mouse sperm DMRs induced by aging, HS and Cd exposure.

Supplemental Table 2. Mouse sperm epigenetic clock model.

